# Dyslexia is characterized by atypical predictive coding specific to the left subcortical auditory pathway

**DOI:** 10.64898/2026.08.17.745222

**Authors:** Heidi Järvikylä, Alejandro Tabas, Katharina von Kriegstein

## Abstract

Developmental dyslexia is a specific, highly prevalent and often debilitating reading and spelling disorder with unknown neurocomputational mechanisms. Here we discovered, in a preregistered functional magnetic resonance imaging study optimized for the subcortical sensory pathway, that dyslexia is characterized by altered predictive coding in left-hemispheric auditory sensory pathway nuclei. The neurocomputational alterations were related to one of the two main dyslexia risk scores, indicating a crucial role for dyslexia pathophysiology.

## Main

Developmental dyslexia (hereafter: dyslexia) is a specific developmental learning disorder characterized by persistent and significant difficulties in acquiring literacy^1^. Dyslexia affects 3-7 % of the population globally^2^ and often impacts educational achievement, mental well-being, and social outcomes^3,4^. Despite the high prevalence and negative impact of dyslexia on the individual and the society, the neurocomputational mechanisms behind key dyslexia symptoms remain unexplained, hampering the development of effective treatment strategies.

Many explanatory theories and models (‘cognitive models’) assume that key dyslexia characteristics result from specific dysfunctions of cognitive abilities such as phonological processing, rapid automatized naming, working memory, and other executive functions. Conversely, other theories of dyslexia emphasize the alterations in slower statistical learning of simple stimuli^5^, or altered adaptation to sensory stimuli both in the auditory and visual modalities^6,7^ (‘sensory models’). One suggestion is that both the cognitive theories and sensory theories can be combined within a predictive coding view of dyslexia^8^. Whether there are predictive coding alterations in dyslexia, however, remains unknown.

The aim of our study was to test whether predictive coding is altered in developmental dyslexia. To do this, we used the auditory subcortical pathway as a model system. We chose this system for two reasons: First, recent studies in typically reading populations showed that the subcortical sensory pathway plays a key role in predictive processing of sounds^9–11^, which is consistent with the massive feedback connections from cortical layer 5/6 to first order sensory thalami^12–14^. Second, there are long-standing unexplained findings of sensory pathway alterations in developmental dyslexia. In the auditory modality these alterations are specific to the left auditory sensory thalamus, i.e., the left medial geniculate body (MGB)^8,15,16^. Surprisingly, these sensory pathway alterations are present for cognitive linguistics tasks, e.g., dyslexics have less left MGB modulation for a speech task in contrast to a control task on the same stimulus input in contrast to typically reading controls. In addition, left MGB alterations are linked to decreased ability to rapidly names letters and numbers (RANln)^8^. RANln is one of the key prognostic factors of dyslexia across orthographies^17,18^ (for a review, see ^19^). Whether left MGB alterations in dyslexia are only related to linguistic processing or in general also to non-speech processing is to date unclear.

To find out whether alterations in left MGB are speech-task specific or rather can be explained by predictive coding mechanism also for non-speech sounds, we tested three preregistered hypotheses^20^. First, we hypothesized that predictive coding in the left MGB is altered in dyslexia even for simple sound stimuli ^14^; second, based on previous research on left MGB involvement, we hypothesized that the potential predictive coding alteration is specific to the left MGB and does not occur in other pathway nuclei such as the IC or the right MGB; and last, that predictive coding alterations in the left MGB are associated with RANln similarly as has been found for speech tasks^8^.

We tested the hypotheses with an experimental paradigm (Figure 1) designed to measure the strength of predictive coding^10^. We used functional magnetic resonance imaging (fMRI) with a sequence optimized for imaging the subcortical sensory pathway nuclei^9^. To test the specificity of the impairment to the left MGB, we measured responses at both left and right MGB and bilateral auditory midbrain (inferior colliculi, IC).

**Figure 1.**
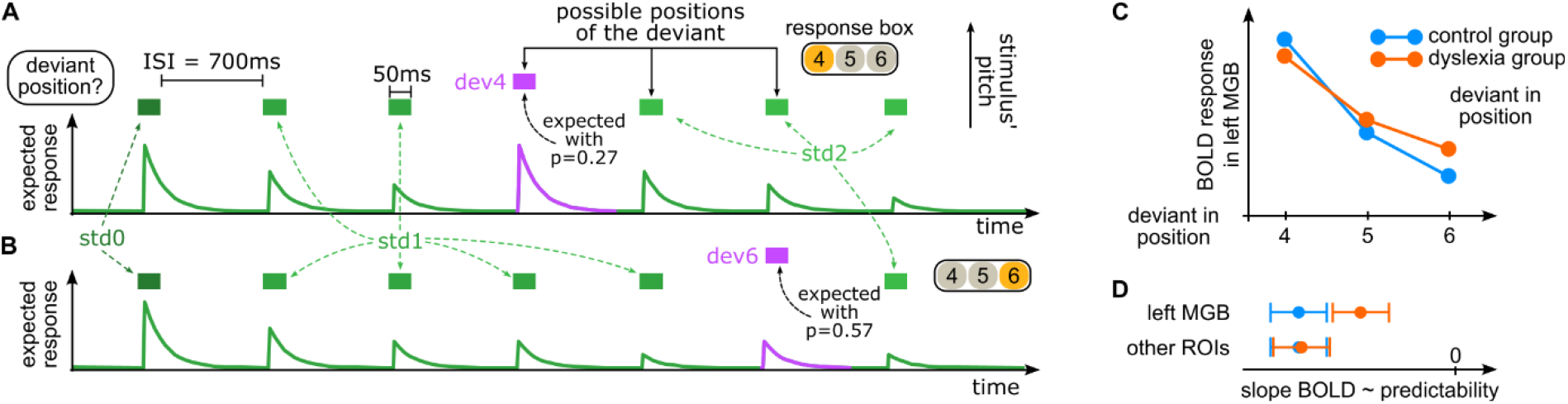
Experimental design and hypotheses. (A) Example trial. A trial is a sequence of 8 pure tones. Participants are instructed to report the location of the tone that differs in pitch (deviant). Deviants occur in 80% of the sequences, in either position 4, 5, or 6. Therefore, after hearing three standards, an ideal observer expects a deviant in position 4 with a probability of ∼26.7%. After hearing four standards, the deviant can only be in positions 5 or 6, and the probability that the trial has a deviant is slightly reduced, resulting in a probability of 36.4% for finding a deviant in position 5. Similarly, after five standards, the probability of a deviant in position 6 is 57.1% (see Methods for details). (B) Hypothesized BOLD responses for the different deviant positions (dev 4, 5, 6). According to predictive coding, responses scale with the predicted probability of the stimuli. Since the onset time of the sequence is unpredictable, we assume higher responses to the first tone of the sequence (dark green; modeled with a specific regressor std0) than to the standard tones preceding (mid green; std1) and following (light green; std2) the deviant (purple; dev4, 5, 6). The standards in position 6 in trials without a deviant (std6) are not displayed in the figure. (C) Preregistered hypotheses for the left MGB. We hypothesized that the dependency of the responses with predictability is reduced in the left MGB in dyslexia. (D) Preregistered hypotheses for the left MGB and other auditory pathway nuclei: The reduction in the predictability of the slope in dyslexia is specific to the left MGB, but does not occur for the other regions of interest. ISI, inter-stimulus interval; dev, deviant; std, standard; BOLD, blood oxygen level dependent, MGB, medial geniculate body, ROI, region of interest.

### Predictive coding is reduced in the left MGB in dyslexia

We first examined whether predictive coding of auditory stimuli differs between individuals with dyslexia and controls in the left MGB (preregistered hypothesis 1). To do this, we fitted a GLM between the BOLD responses and the experimental conditions (*std0, std1, std2, dev4, dev5, dev6, std6*; Figure 1) and then performed two second level analyses.

In the first second level analyses, BOLD responses in the left MGB of the control (β = -84.34, SE = 12.10, t = −6.97, p = 3.18 × 10^-12^), and the dyslexia (β = -68.70, SE = 12.01, p = 2.64 x 10^-7^) group were strongly correlated with predictability. This result is in agreement with previous findings in typical readers^9–11^. The results were obtained by fitting a linear mixed-effects model (LMM) regressing voxel-wise estimates in left MGB to each condition to their predictability per each of the two groups. The dependency of the responses with predictability was measured as the value of that regressor β. The models included random effects for subject (intercepts and slopes) and run (intercepts).

In agreement with our first preregistered hypothesis, the dependency with predictability in the left MGB was significantly weaker in the dyslexia than in the control group (β = 17.1, SE = 3.32, p = 2.64 x 10^-7^, Cohen’s d = 0.44). This result was based on an LMM including both groups and including an interaction term between predictability and group. The strength of the effect was measured as the beta value of the regressor of the interaction term. The results could not be attributed to behavioral differences between groups, because they were comparable (largest Cohen’s d = 0.21 for all behavioral effects, see Methods).

### Predictive Coding alterations are not specific to left MGB but also encompass the left auditory midbrain (IC)

To test our second preregistered hypothesis, we evaluated the specificity of the response alterations to left MGB. To do this, we fitted LMMs including the interaction between group and predictability for each of the three remaining subcortical ROIs (MGB-R, IC-L, and IC-R; Figure 2). In congruence with our hypothesis, the right-lateralized nuclei, for the right MGB and right IC, the dependency of responses on predictability was similar across the two groups, with negligible interaction (Cohen’s d between -0.01 and -0.03; Figure 2).

**Figure 2.**
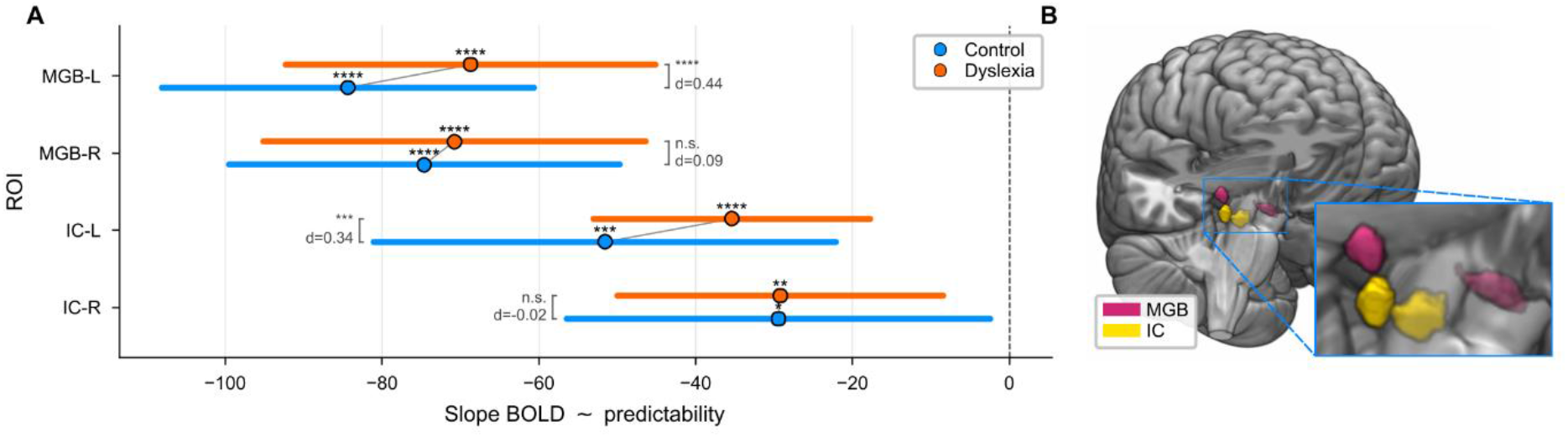
MRI results reflecting reduced predictive coding in left auditory thalamus (MGB-L) and midbrain (IC-L). The plot shows coefficient estimates (± 95% CI) for the dependency of response strength with predictability in the dyslexia (orange) and control (blue) groups. From top to bottom, rows show estimates for the left MGB, right MGB, left IC, and right IC. More negative values indicate greater effects of predictability and thus stronger predictive coding. The prediction slope was significantly weaker in the dyslexic group than in controls in the left MGB (p = 2.64 x 10^-7^, d = 0.44) and the left IC (p = 0.0002, d = 0.34), reflecting reduced predictive coding. This group difference was specific to the left hemisphere and did not occur in the right-hemispheric nuclei (MGB-R: d = 0.09; IC-R: d = -0.02). Cohen’s ds were computed as the ratio between the mean estimate difference and the pooled variance, as estimated form the random slopes. Significance levels: * p < 0.05, ** p < 0.01, **** p < 0.0001; ns = not significant. N= 24 in dyslexia group N=24 in control group. MGB, medial geniculate body; IC, inferior colliculus.

**Figure 3.**
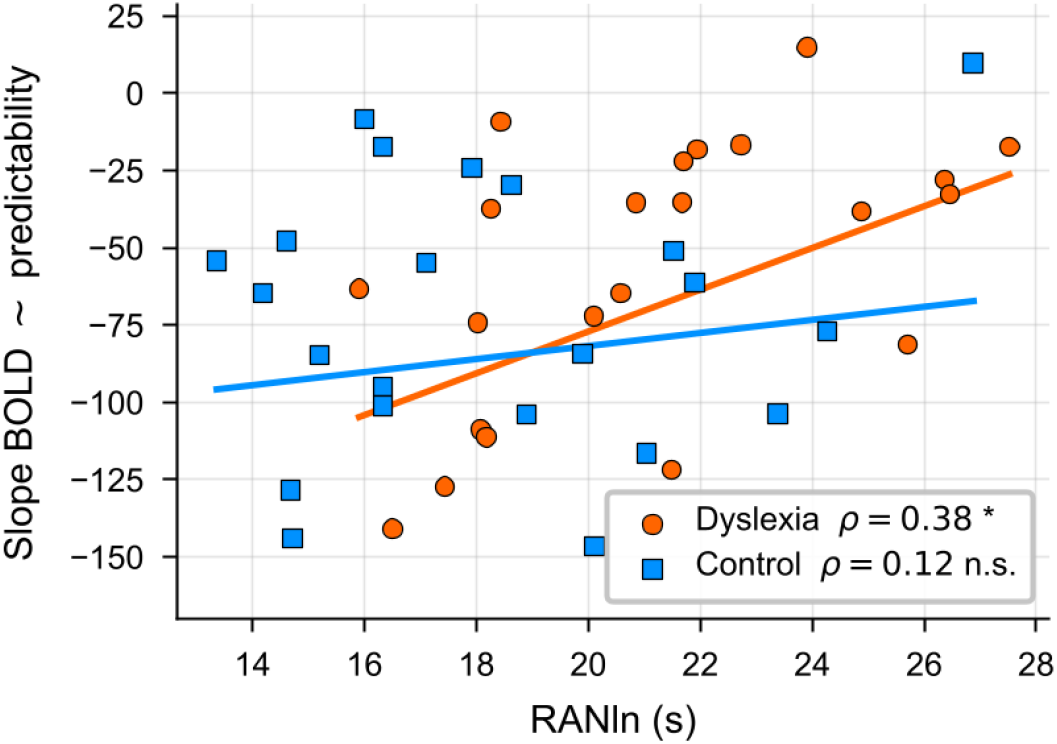
Correlation between key diagnostic score (RANln) and prediction slope in the left MGB in dyslexia and pair-wise matched controls. There was a significant correlation between the alteration in predictive coding in the left MGB in dyslexia and the time required to rapidly name letters and numbers (r = 0.38, p = 0.033), while there was no significant correlation between the variables in controls (r = 0.12, p = 0.29).

However, contrary to our hypothesis, the left IC has similar response alterations to the left MGB in dyslexia: the dependency on predictability in the left IC was also significantly weaker in dyslexia than in the control group (β = 15.1, SE = 4.03, p = 0.0002, Cohen’s d = 0.34). These differences in the results between left MGB and left IC, and the right MGB and right IC cannot be attributed to differences in statistical power as all four ROIs did not substantially differ in their temporal signal-to-noise ratios (tSNRs) across groups (see Methods).

### Reduced Predictive Coding in left MGB Linked to key Diagnostic Score

To test our third hypothesis, we ran one-tailed Pearson’s correlations between the prediction slope in left MGB and RANln time for the dyslexic participants. In agreement with the hypothesis, RANIn performance was significantly correlated to the prediction slope in left MGB in dyslexic participants (*r* = 0.38, *p* = 0.035). Due to the surprising BOLD response alterations found in the left IC in dyslexia for predictability, we also ran an exploratory correlation between the prediction slope in the left IC and RANln time for the dyslexic participants. The RANln performance was significantly correlated to the prediction slope also in the left IC (*r* = 0.39, *p* = 0.03).

### Predictive coding – a computational mechanism for linking sensory and linguistic alterations in dyslexia

Our study revealed predictive coding alterations in developmental dyslexia that were specific to two auditory sensory pathway nuclei – the left MGB and left IC. The alterations were present during the processing of simple auditory stimuli, and directly linked to a key behavioral linguistic dyslexia measure, i.e., the ability to rapidly name letters and numbers. This is a strong indication that predictive coding alterations in left-hemispheric subcortical sensory pathway nuclei might be a direct link between ‘cognitive models’ and ‘sensory models’ as explanation for dyslexia symptoms.

Competing computational frameworks have been introduced to explain altered neural processing in dyslexia. One such explanation is faster decay of adaptation, attempting to account for reduced adaptation to repeated stimuli across categories^6,7^. Another one, the decreased use of recent stimulus statistics^22,23^, addresses impaired strengthening of standard representations^7^. Both adaptation models can be reconciled by the predictive coding framework^8,24^. Reduced predictive coding leads to higher prediction errors despite the repetition of a stimulus, manifesting in reduced adaptation. Similarly, the impaired strengthening of stimulus statistics can be explained with atypical predictions^7^. Moreover, while the adaptation models reflect a phenomenological description of stimulus-driven response reduction, the predictive coding framework offers a parsimonious computational explanation that listeners build and update an internal generative model of the environment, linking adaptation directly to a specific sensory processing strategy. Critically, the predictability-by-group interaction in the present study established that individuals with dyslexia do not employ predictions as efficiently as controls, a result that goes beyond what adaptation experiments could establish.

Also the often observed difficulty in dyslexia with speech processing, whether at the level of phonemes^25^ or speech in noisy environments^26^, is in line with altered predictive coding mechanism in dyslexia. The difficulties with accurately predicting fast, complex, and highly predictable stimuli such as speech sounds in dyslexia would lead to slower and less accurate phoneme recognition, and on higher levels of the computational hierarchy, words and utterances. In accordance with hierarchical models of predictive coding^13,27,28^, the lower, subcortical levels, e.g., the MGB, process faster dynamics making them essential for accurate tracking of speech sounds^8,29^.

Whether the reduced predictive coding in the left MGB reflects intrinsic subcortical dysfunction or diminished top-down corticofugal drive remains unresolved. Structural connectivity between the left MGB and motion-sensitive planum temporale is selectively reduced in dyslexia and correlates with RANln^32^, raising the possibility that atypical corticothalamic input may contribute to the observed left-hemispheric sensory pathway alterations. However, the alterations might also originate within the left MGB itself, as post-mortem histological results show left MGB alterations^16^. Whether such histological alterations are also present in IC, is unknown.

The left-hemispheric specificity of the predictive coding alterations in dyslexia is consistent with the broader literature: the left auditory thalamus is preferentially implicated in processing rapidly time-varying acoustic features critical for phonological representation^29,34^, and its modulation as well as IC modulation is behaviorally relevant during speech recognition in typically reading participants^29^. The left-hemispheric bias also aligns with the auditory temporal processing^35^ and phonological difficulties^36,37^ widely documented in dyslexia. The left-hemispheric alterations also mirror findings in the visual domain where LGN hemispheric asymmetry was associated with RANln performance in males with dyslexia^33^.

Our study shows that the consistently found relationship between left hemispheric sensory thalamic alterations in dyslexia and naming speed^8,33,38^ generalizes to subcortical predictive coding measures. Together, the findings directly link sensory thalamus to a key behavioral measure that predicts reading ability^17,39^. Whether this is causally antecedent to dyslexia or exacerbated throughout reading life requires further investigation.

Dyslexia is a multifactorial disorder with a heterogeneity of symptoms across individuals^40^ and it might therefore not be possible to explain dyslexia as one disorder. Despite this, the importance of finding a relationship with one of the two main cognitive scores that predict developmental dyslexia^41,42^ helps move the field further in explaining the mechanics of the disorder.

In sum, our results provide first evidence of left-lateralized thalamic and midbrain dysfunction of predictive coding in dyslexia. This subcortical alteration situates dyslexia within a predictive coding framework, leading to an understanding of the computational alterations in the disorder.

## Methods

This study was approved by the Ethics Committee of TUD Dresden University of Technology, Germany (EK 315062019). All participants gave their written informed consent before data acquisition and were remunerated for their participation in the study financially or in course credits.

### Participants

Forty-eight adult native German speakers participated in the study; one group included 24 participants with dyslexia and the other 24 typically reading controls (hereafter: controls). We determined the sample size based on the results of a pilot study with 7 participants with dyslexia and 7 control participants. The calculations, using G*Power (v 3.1.9.7; ^43^) indicated a sample of N > 10.2 is required for a statistical power of 0.8 for testing hypotheses 1 and 2, and *n* = 24 for a power of 0.9 for the correlation in hypothesis 3. None of the participants reported a history of psychiatric or neurological disorders in the previous two years, current use of psychoactive medications, or hearing difficulties. Normal hearing was confirmed with pure tone audiometry (250–8000 Hz; Inventis Piano Plus, Padova, Italy, https://www.inventis.it) with a threshold equal or below 25 dB in the frequency range used in the task, 1300–1600 Hz.

Thirteen participants in the dyslexia group had a formal, written diagnosis, the other 11 reported severe reading and spelling difficulties since childhood. Group allocation was confirmed with 22 of the dyslexia participants scoring at least 1.5 SD below control mean on two or more literacy measures. Of the 2 others, one had an official diagnosis, and the other scored 1.5 SD below control mean in one, and close to 1.5 SD below control mean in two other measures. Each participant in the control group was pair-wise matched with a participant in the dyslexia group in age, sex, handedness (assessed using the Short Form of the Edinburg Handedness Inventory ^44^), educational level, and non-verbal intelligence quotient (IQ). IQ was assessed using the German adapted version of the Wechsler Adult Intelligence Scale (WAIS-IV; Wechsler, 2012; German version edited by ^45^) confirming that participants’ intellectual abilities were in the normal range (score ≥85).

All participants performed a formal assessment on reading accuracy and comprehension (LGVT-5-12+; ^46^), fluency (SLRT II; ^47^ and spelling (RST-ARR; ^48^), and rapid automatized naming of letters and numbers (i.e. RANln; ^49^, German protocol in ^8^) and phonological manipulation (Spoonerism; in-house test, unpublished, used in ^8^).

Participants were also screened for autism spectrum disorders (Autism Spectrum Quotient; ^50^; German version adapted from ^51^), adult attention-deficit/hyperactivity disorder (ASRS Screener v1.1; ^52^), and prosopagnosia (PI20; ^53^).

**Table 1.**
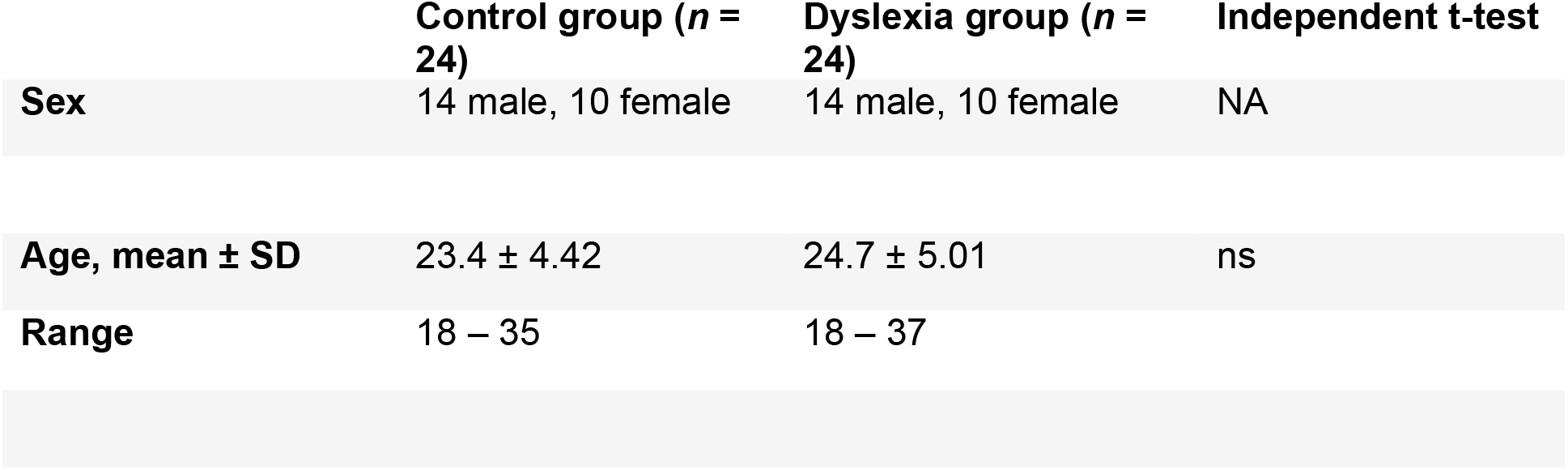
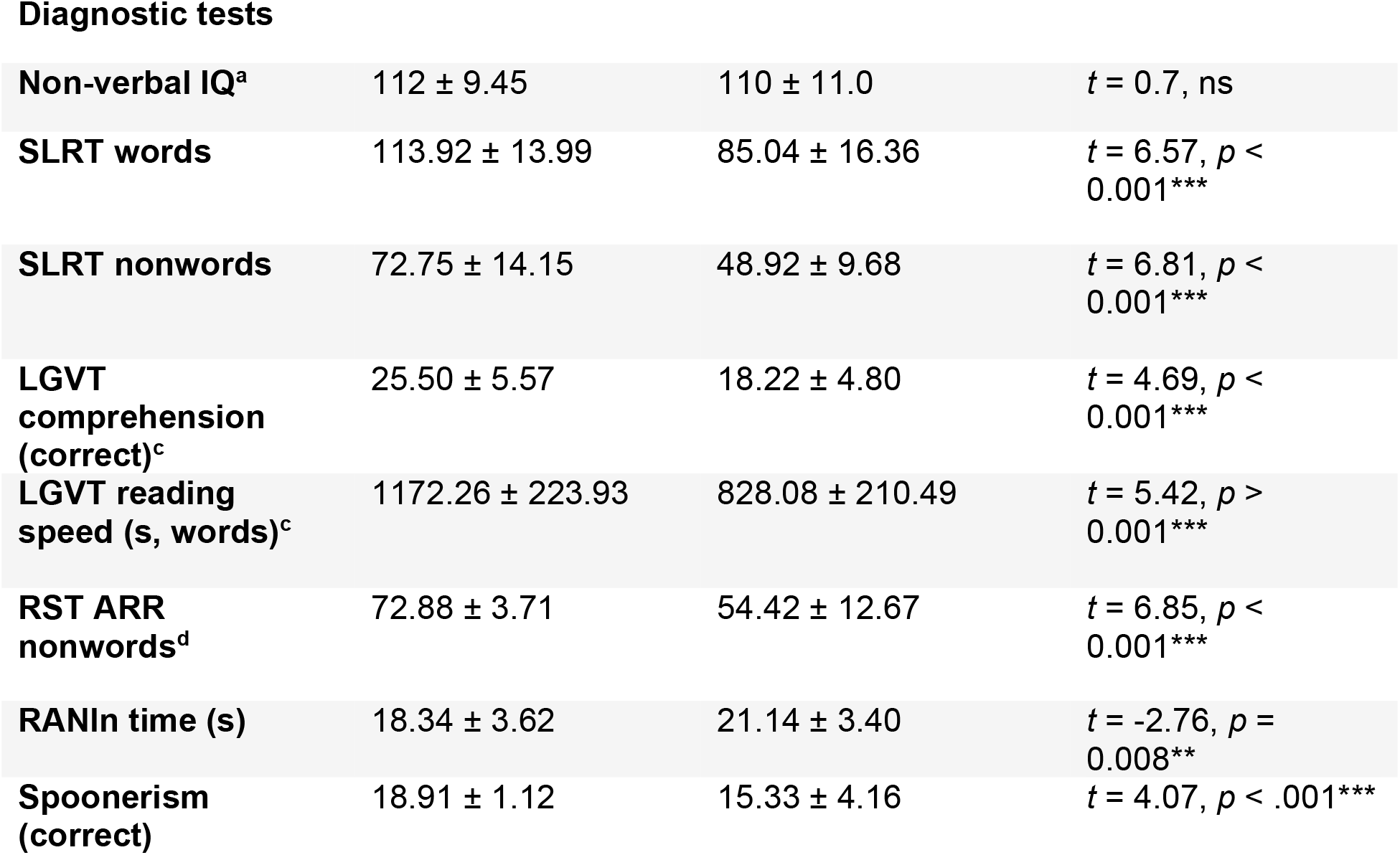
Demographic and cognitive measures (mean ± SD) of 24 control and 24 dyslexic participants. Participants in the control group and dyslexia group were pairwise matched in chronological age, gender, non-verbal intelligence quotient (IQ), and handedness. ^a^ WAIS = The Perceptual Reasoning Index in the Wechsler Adult Intelligence Scale IV ^45^, M = 100; SD = 10 ^b^ SLRT II = the second edition of the Salzburg reading and spelling tests ^47^ ^c^ LGVT 5-12+ = Reading speed and comprehension test for grades 5-12+; 2nd, extended and newly standardized edition ^46^ ^d^ RST-ARR = Rechtschreibtest – Aktuelle Rechtschreibregelung; 3rd, new standardized edition ^48^

### Experimental paradigm

The stimuli and design were adapted from ^10^. The stimuli were pure tones of 1300 Hz, 1400 Hz, and 1600 Hz, each 50 ms in duration including 5 ms in/out ramps. The pure tones were combined into 6 standard-deviant combinations, all of which were used as the standard and the deviant. The differences between the two tones (Δf) were 100 Hz, 200 Hz and 300 Hz, and were all well distinguishable by typical readers as well as participants with dyslexia ^10,54^. The six standard-deviant combinations were each played in a separate block so that within the block there was a strong expectation on the standard and deviant frequencies.

Each trial within a block consisted of a sequence of eight tones with a 700 ms inter-stimulus interval (ISI). The inter-trial interval (ITI) was jittered so that the deviants were separated by 5000 ms on average, with a minimum and maximum ITI of 1500 ms and 11 000 ms, respectively. Seven of the pure tones were repetitions of the same frequency (standard) and one tone deviated in frequency (deviant). The deviant occurred in the sequence at position 4, 5, or 6 (Figure 1). Participants were asked to report, via a button press, the position of the deviant in each sequence as quickly and as accurately as possible. In 20% of the trials there was no deviant, i.e., the standard tone was repeated eight times. We manipulated the participants’ expectations independently of stimulus regularity using two abstract rules that were disclosed to the participants: 1) 80% of the sequences contain one deviant, the remaining 20% contain no deviant; 2) the deviant can occur in positions four, five or six within the sequence. Thus, after hearing three standards, an ideal observer expects a deviant in position four with a probability of ca. 26.7%. After hearing four standards, the deviant can only be in positions 5 or 6, and the probability that the trial has a deviant is slightly reduced, resulting in a probability of 36.4% for finding a deviant in position 5. Similarly, after five standards, the probability of a deviant in position 6 is 57.1%. Thus, although deviants are equally likely in all positions based on the statistical history, participants held different priors on the likelihood of hearing a deviant across deviant positions.

The experiment consisted of 12 runs of the same task, obtained in three separate fMRI sessions. Each run contained 6 blocks of 10 trials, and lasted approximately 12 minutes each. The trials within a block used one of the six possible combinations of pure tones, resulting in each sequence within a block to have the same standard and deviant. This way, within a block, the deviant and standard frequencies were known to the participant, while the position of the deviant was unknown from trial to trial. The order of the blocks within the experiment was randomized. The position of the deviant was pseudorandomized across all trials of a run, leading each deviant position to occur exactly 20 times per run, but an unknown number of times within a block. Adding this constraint made it possible for us to maintain the same *a priori* probability for all deviant positions in each block. Additionally, 23 silent gaps of 5300 ms (i.e., null events with the same duration as the tone sequences) were randomly located in each run ^55^.

Each run was separated by a minimum one-minute break, during which the participants could rest. Fieldmaps and a whole-head EPI were acquired between the first and second run. During the first session, the anatomical image was acquired between the second and third run.

### Data acquisition

MRI data were acquired using a 3T Tim Trio (Siemens Healthcare GmbH, Erlangen, Germany), and a Siemens Magnetom Prisma 3 Tesla machine (Siemens Healthineers, Erlangen, Germany) with a 32-channel head coil. Each pairwise-matched dyslexia-control pair was measured using the same MRI machine. We opted for the 32-channel head-coil as it allowed us to fit participants with larger head-sizes in the coil, despite the space needed also for the headphones. The data were acquired with two different setups due to a system upgrade during the data acquisition phase of the study.

Functional MRI data were acquired using an echo planar imaging (EPI) sequence with a partial coverage with 24 slices of the slab. The EPI sequence had the following acquisition parameters: TR = 1900ms, TE = 42ms, flip angle 66°, matrix size 88 × 88, field of view (FoV) 154mm×154mm, voxel size 1.75mm or 1.8mm isotropic (Tim Trio, and Magnetom Prisma, respectively), bandwidth per pixel 1386Hz/px, and interleaved acquisition. The slab was oriented in parallel to the superior temporal gyrus so that the slices encompass the IC, the MGB and the superior temporal gyrus.

Structural data were acquired using MPRAGE T1 protocol with 1 mm isotropic resolution, TE = 1.95 ms or 1.97 ms (in Tim Trio and Prisma, respectively), TR = 2000 ms, IT = 880 ms, flip angle = 8°.

Participants’ heart rate and respiratory rate were measured by means of a pulse oximeter. Due to technical reasons, there is no physio data for all sessions. Fourteen participants in the dyslexia group have physio for one or more sessions, 13 controls have it for one or more sessions.

The stimuli were presented using MATLAB (Matlab R2019b), with the PsychoPy2 (1.90.3) and Psychophysics Toolbox extensions and delivered through OptoACTIVE Active Noise Control system with OptoActive II headphones (Optoacoustics Ltd.). Loudness was adjusted individually for each participant to a comfortable hearing level prior to starting the data acquisition. Participants received visual feedback after every trial for a maximum duration of 2000 ms indicating whether they had responded correctly to each trial.

### Data preprocessing

The data were preprocessed using a pipeline coded in Nipype, version 1.8.6 ^56^, executed using python v 3.11.6, IPython 8.16.1, the Statistical Parametric Mapping toolbox, version 12 (SPM12; ^57^); Freesurfer, version 7.3.2 ^58^; the MRIB Software Library ^59–62^; and the Advanced Normalization Tools, version 2.5.1 (ANTS; ^63^). All data were coregistered to the Montreal Neurological Institute (MNI) MNI152 template with 1 mm isotropic resolution (ICBM 152, version 2009; ^64,65^).

First, we processed the structural data. We used the reconall-routine in Freesurfer to calculate the boundaries between gray and white matter for later registering the functional data to the structural images, and ANTS to compute the transformation between the structural data and the MNI152 template.

Next, we realigned the functional runs. For that, we used the *FieldMap Toolbox* in SPM to calculate the geometric distortions in the EPI images caused by field inhomogeneities. After that, we used the *Realign* in SPM to execute motion and distortion correction. Motion artifacts were calculated using *ArtifactDetect* in SPM, and added to the design matrix later when estimating the BOLD responses.

Then, we co-registered the functional data to the MNI152 space. We used *BBregister* in Freesurfer to calculate the transformation between the functional data and the structural image, using the boundaries between gray and white matter of the structural data and the whole-brain EPI as an intermediate step. The final transformation was computed as the concatenation of the functional-to-structural and structural-to-MNI transformations, and was applied using ANTS. As the resolution of the MNI space (1 mm isotropic) was higher than the resolution of the functional data (1.75 and 1.8 mm isotropic), the transformation led to spatial oversampling. Before running the linear models, the data were resampled to native resolution.

Finally, the co-registered data were smoothed using a 2 mm full-width half-maximum kernel Gaussian kernel with the *Smooth* function in SPM.

### Estimation of the BOLD responses

First and second level analyses were coded in Nipype and executed using SPM. The statistical analyses of the model estimations in the ROIs were performed using custom MATLAB code (Matlab R2023b, v 23.2).

The design matrix of the first level analysis included seven conditions: first standard (std0), standards before the deviant (std1), standards after the deviant (std2), deviants 4, 5, and 6 (dev4, dev5, and dev6, respectively), and standard in position 6 in trials without a deviant (std6; Figure 1).

We modeled conditions std1 and std2 using linear parametric modulation ^66^, with their linear factors coded for the position of the sound within the sequence. The first standard was modeled separately from the following standards preceding the deviant in order to contrast the responses to the first and the adapted standards to locate voxels that exhibit adaption. Similarly, we modeled the standards occurring before and after the deviant separately as designing a set of linear factors that are simultaneously valid for both std1 and std2 would require making assumptions on any possible recovery experienced during the presentation of the deviant. In addition to these main regressors, the design matrix also included the physiological PhysIO where available, and artefact regressors of no-interest.

### Definition of the ROIs

To locate the voxels corresponding to our regions of interest, the left and right IC, and the left and right MGB, we used an anatomical atlas of the subcortical auditory pathway ^67^. The atlas was compiled using histological data from the BigBrain project, postmortem MRI data, as well as in-vivo fMRI data in response to natural sounds. The masks used in the current study were computed based on the latter, the fMRI data, due to its similarity to the current experimental setup.

### Temporal Signal-to-Noise Ratio of the ROIs

The four ROIs did not substantially differ in their temporal signal-to-noise ratios (tSNRs) across groups (all mean tSNR values were between 25.81 and 27.29; all standard deviations were between 5.56 and 7.51). All pairwise comparisons were non-significant (Cohen’s d between - 0.04 and 0.21) except for control IC-L > control MGB-L (pFDR = 0.02, Cohen’s d = 0.21), and control IC-L > control MGB-R (pFDR = 0.04, Cohen’s d = 0.17). Thus, differences in the results cannot be attributed to differences in statistical power.

### Behavioral task results

The difference in the left MGB activation between controls and participants with dyslexia cannot be explained by behavioral differences. Both groups showed high performance to all deviant positions (controls: 98 ± 4%, 98 ± 4%, and 99 ± 4%, mean accuracies ± standard error of the mean, for deviants in positions 4, 5, and 6, respectively; participants with dyslexia: 98 ± 4%, 99 ± 2%, and 99 ± 2%) indicating that participants were attentive and both groups could behaviorally perform the task with ease. Reaction times (controls: *RT* = 618 ± 19 ms, 571 ± 17 ms, and 476 ± 15 ms; participants with dyslexia: *RT* = 622 ± 21 ms, 554 ± 18 ms, 465 ± 16 ms) in both groups were shorter for the more expected deviants, indicating a behavioral benefit of predictability to both controls as well as participants with dyslexia. This was further confirmed with Wilcoxon rank-sum tests for each of the six comparisons, all *p*s > 0.05. RTs were significantly shorter for deviants at position six than for deviants at positions 4 and 5 for controls (Cohen’s *d* = −1.5, and *d* = −1.7, respectively; *p* < 0.000) and for readers with dyslexia (Cohen’s *d* = −1.7, and *d* = −1.3, respectively; *p* < 0.000). They were also shorter for deviants at position 5 than deviants at position 4 in controls (Cohen’s *d* = −0.7, *p* < 0.000) and readers with dyslexia (Cohen’s *d* = −0.8, *p* < 0.000). Statistical difference was assessed separately with two-tailed Ranksum tests with *n* = 24 in each group, Holm-Bonferroni corrected for three comparisons).

## Data Availability

All code used for data processing and analysis are publicly available and the study was preregistered and is available at https://osf.io/dx6za.

## References

1. World Health Organization. The ICD-10 Classification of Mental and Behavioural Disorders. ICD-11 for Mortality and Morbidity Statistics vol. 55 135–139 https://icd.who.int/browse11/l-m/en#/ http://id.who.int/icd/entity/1008636089 (2025).

2. Wagner, R. K. et al. The Prevalence of Dyslexia: A New Approach to Its Estimation. J. Learn. Disabil. 53, 354–365 (2020).

3. Cassidy, L., Reggio, K., Shaywitz, B. A., Holahan, J. M. & Shaywitz, S. E. Correctional Education Association Dyslexia in Incarcerated Men and Women. Source J. Correct. Educ. 72, 61–81 (1974).

4. Morte-Soriano, M. R. & Soriano-Ferrer, M. Beyond Reading: Psychological and Mental Health Needs in Adolescents with Dyslexia. Pediatr. Rep. 16, 880–891 (2024).

5. Ballan, R., Durrant, S. J., Manoach, D. S. & Gabay, Y. Failure to consolidate statistical learning in developmental dyslexia. Psychon. Bull. Rev. 30, 160–173 (2023).

6. Perrachione, T. K. et al. Dysfunction of Rapid Neural Adaptation in Dyslexia. Neuron 92, 1383–1397 (2016).

7. Beach, S. D. et al. The Neural Representation of a Repeated Standard Stimulus in Dyslexia. Front. Hum. Neurosci. 16, (2022).

8. Díaz, B., Hintz, F., Kiebel, S. J. & Kriegstein, K. V. Dysfunction of the auditory thalamus in developmental dyslexia. Proc. Natl. Acad. Sci. U. S. A. 109, 13841–13846 (2012).

9. Tabas, A., Kiebel, S., Marxen, M. & von Kriegstein, K. Fast frequency modulation is encoded according to the listener expectations in the human subcortical auditory pathway. Imaging Neurosci. 2, imag–2–00292 (2024).

10. Tabas, A., Mihai, G., Kiebel, S., Trampel, R. & Kriegstein, K. V. Abstract rules drive adaptation in the subcortical sensory pathway. eLife 9, 1–19 (2020).

11. Ara, A., Provias, V., Sitek, K., Coffey, E. B. J. & Zatorre, R. J. Cortical–subcortical interactions underlie processing of auditory predictions measured with 7T fMRI. Cereb. Cortex N. Y. NY 34, bhae316 (2024).

12. Sillito, A. M. & Jones, H. E. Corticothalamic interactions in the transfer of visual information. Philos. Trans. R. Soc. Lond. B. Biol. Sci. 357, 1739–1752 (2002).

13. Asilador, A. & Llano, D. A. Top-Down Inference in the Auditory System: Potential Roles for Corticofugal Projections. Front. Neural Circuits 14, (2021).

14. Antunes, F. M. & Malmierca, M. S. Corticothalamic Pathways in Auditory Processing: Recent Advances and Insights From Other Sensory Systems. Front. Neural Circuits 15, (2021).

15. Tschentscher, N., Ruisinger, A., Blank, H., Díaz, B. & von Kriegstein, K. Reduced Structural Connectivity Between Left Auditory Thalamus and the Motion-Sensitive Planum Temporale in Developmental Dyslexia. J. Neurosci. 39, 1720–1732 (2019).

16. Galaburda, A. M., Menard, M. T. & Rosen, G. D. Evidence for aberrant auditory anatomy in developmental dyslexia. Proc. Natl. Acad. Sci. U. S. A. 91, 8010–8013 (1994).

17. Landerl, K. et al. Predictors of developmental dyslexia in European orthographies with varying complexity. J. Child Psychol. Psychiatry 54, 686–694 (2013).

18. Caravolas, M. et al. Common Patterns of Prediction of Literacy Development in Different Alphabetic Orthographies. Psychol. Sci. 23, 678–686 (2012).

19. Norton, E. S. & Wolf, M. Rapid Automatized Naming (RAN) and Reading Fluency: Implications for Understanding and Treatment of Reading Disabilities. Annu. Rev. Psychol. 63, 427–452 (2012).

20. Järvikylä, H., Tabas, A. & Von Kriegstein, K. Aberrant predictive coding in subcortical sensory pathways in developmental dyslexia. (2025).

21. Benjamini, Y. & Hochberg, Y. Controlling the False Discovery Rate: A Practical and Powerful Approach to Multiple Testing. J. R. Stat. Soc. Ser. B Methodol. 57, 289–300 (1995).

22. Jaffe-Dax, S., Frenkel, O. & Ahissar, M. Dyslexics’ faster decay of implicit memory for sounds and words is manifested in their shorter neural adaptation. eLife 6, e20557 (2017).

23. Jaffe-Dax, S., Lieder, I., Biron, T. & Ahissar, M. Dyslexics’ usage of visual priors is impaired. J. Vis. 16, 10 (2016).

24. Jaffe-Dax, S., Raviv, O., Jacoby, N., Loewenstein, Y. & Ahissar, M. A Computational Model of Implicit Memory Captures Dyslexics’ Perceptual Deficits. J. Neurosci. 35, 12116–12126 (2015).

25. Snowling, M. Dyslexia as a Phonological Deficit: Evidence and Implications. Child Psychol. Psychiatry Rev. 3, 4–11 (1998).

26. Ziegler, J. C., Pech-Georgel, C., George, F. & Lorenzi, C. Speech-perception-in-noise deficits in dyslexia. Dev. Sci. 12, 732–745 (2009).

27. Carbajal, G. V. & Malmierca, M. S. The Neuronal Basis of Predictive Coding Along the Auditory Pathway: From the Subcortical Roots to Cortical Deviance Detection. Trends Hear. 22, 2331216518784822 (2018).

28. Huang, Y. T. et al. Crossmodal hierarchical predictive coding for audiovisual sequences in the human brain. Commun. Biol. 7, 965 (2024).

29. von Kriegstein, K., Patterson, R. D. & Griffiths, T. D. Task-Dependent Modulation of Medial Geniculate Body Is Behaviorally Relevant for Speech Recognition. Curr. Biol. 18, 1855–1859 (2008).

30. Antunes, F. M. & Malmierca, M. S. An Overview of Stimulus-Specific Adaptation in the Auditory Thalamus. Brain Topogr. 27, 480–499 (2014).

31. Ayala, Y. A. & Malmierca, M. S. Stimulus-specific adaptation and deviance detection in the inferior colliculus. Front. Neural Circuits 6, 1–16 (2013).

32. Tschentscher, N., Ruisinger, A., Blank, H., Díaz, B. & Kriegstein, K. von. Reduced Structural Connectivity Between Left Auditory Thalamus and the Motion-Sensitive Planum Temporale in Developmental Dyslexia. J. Neurosci. 39, 1720–1732 (2019).

33. Müller-Axt, C., Kauffmann, L., Eichner, C. & von Kriegstein, K. Dysfunction of the magnocellular subdivision of the visual thalamus in developmental dyslexia. Brain 148, 252–261 (2025).

34. Mihai, P. G. et al. Modulation of tonotopic ventral medial geniculate body is behaviorally relevant for speech recognition. eLife 8, e44837 (2019).

35. Marchesotti, S. et al. Selective enhancement of low-gamma activity by tACS improves phonemic processing and reading accuracy in dyslexia. PLoS Biol. 18, e3000833 (2020).

36. Temple, E. et al. Disruption of the neural response to rapid acoustic stimuli in dyslexia: Evidence from functional MRI. Proc. Natl. Acad. Sci. U. S. A. 97, 13907–13912 (2000).

37. Landi, N., Mencl, W. E., Frost, S. J., Sandak, R. & Pugh, K. R. An fMRI study of multimodal semantic and phonological processing in reading disabled adolescents. Ann. Dyslexia 60, 102–121 (2010).

38. Müller-Axt, C., Anwander, A. & Kriegstein, K. von. Altered Structural Connectivity of the Left Visual Thalamus in Developmental Dyslexia. Curr. Biol. 27, 3692–3698.e4 (2017).

39. Wolf, M. & Bowers, P. G. The double-deficit hypothesis for the developmental dyslexias. J. Educ. Psychol. 91, 415–438 (1999).

40. Peterson, R. L. & Pennington, B. F. Developmental Dyslexia. Annu. Rev. Clin. Psychol. 11, 283–307 (2015).

41. Carioti, D., Masia, M. F., Travellini, S. & Berlingeri, M. Orthographic depth and developmental dyslexia: a meta-analytic study. Ann. Dyslexia 71, 399–438 (2021).

42. McWeeny, S. et al. Rapid Automatized Naming (RAN) as a Kindergarten Predictor of Future Reading in English: A Systematic Review and Meta-analysis. Read. Res. Q. 57, 1187–1211 (2022).

43. Faul, F., Erdfelder, E.Lang, A.-G. & Buchner, A. G*Power 3: a flexible statistical power analysis program for the social, behavioral, and biomedical sciences. Behav. Res. Methods 39, 175–191 (2007).

44. Veale, J. F. Edinburgh Handedness Inventory - Short Form: A revised version based on confirmatory factor analysis. Laterality 19, 164–177 (2014).

45. Petermann, F. WAIS-IV. Wechsler Adult Intelligence Scale – Fourth Edition. Deutschsprachige Adaptation Der WAIS-IV von D. Wechsler. (Pearson Assessment, Frankfurt a. M., 2012).

46. Schneider, W., Schlagmüller, M. & Ennemoser, M. LGVT 5-12 +. Lesegeschwindigkeits-Und –Verständnistest Für Die Klassen 5–12. 2. Erweiterte Und Neu Normierte Auflage. (Hogrefe, Göttingen, 2017).

47. Moll, K. & Landerl, K. SLRT-II. Lese-Und Rechtschreibtest. Weiterentwicklung Des Salzburger Lese-Und Rechtschreibtests. (Huber, Bern, 2010).

48. Ibrahimovic, N. & Bulheller, S. Rechtschreibtest – Aktuelle Rechtschreibregelung (RST-ARR). (Pearson, Frankfurt a. M., 2013).

49. Denckla, M. B. & Rudel, R. G. Rapid ‘automatized’ naming (R.A.N.): Dyslexia differentiated from other learning disabilities. Neuropsychologia 14, 471–479 (1976).

50. Baron-Cohen, S., Wheelwright, S., Skinner, R., Martin, J. & Clubley, E. The Autism-Spectrum Quotient (AQ): Evidence from … J. Autism Dev. Disord. 31, 5–17 (2001).

51. Freitag, C. M. et al. Evaluation der deutschen Version des Autismus-Spektrum-Quotienten (AQ) - die Kurzversion AQ-k. Z. Für Klin. Psychol. Psychother. 36, 280–289 (2007).

52. Kessler, R. C. et al. The World Health Organization adult ADHD self-report scale (ASRS): a short screening scale for use in the general population. Psychol. Med. 10.1017/s0033291704002892 (2005) doi:10.1017/s0033291704002892.

53. Shah, P., Gaule, A., Sowden, S., Bird, G. & Cook, R. The 20-item prosopagnosia index (PI20): a self-report instrument for identifying developmental prosopagnosia. R. Soc. Open Sci. 2, 140343 (2015).

54. Witton, C., Swoboda, K., Shapiro, L. R. & Talcott, J. B. Auditory frequency discrimination in developmental dyslexia: A meta-analysis. Dyslexia 26, 36–51 (2020).

55. Friston, K. J., Zarahn, E., Josephs, O., Henson, R. N. A. & Dale, A. M. Stochastic Designs in Event-Related fMRI. NeuroImage 10, 607–619 (1999).

56. Gorgolewski, K. et al. Nipype: A flexible, lightweight and extensible neuroimaging data processing framework in Python. Front. Neuroinformatics 5, (2011).

57. SPM12: Statistical Parametric Mapping. (2020).

58. Fischl, B. FreeSurfer. NeuroImage vol. 62 774–781 (2012).

59. Greve, D. N. & Fischl, B. Accurate and Robust Brain Image Alignment using Boundary-based Registration. NeuroImage 48, 63–72 (2009).

60. Jenkinson, M. & Smith, S. A global optimisation method for robust affine registration of brain images. Med. Image Anal. 5, 143–156 (2001).

61. Jenkinson, M., Bannister, P., Brady, M. & Smith, S. Improved Optimization for the Robust and Accurate Linear Registration and Motion Correction of Brain Images. NeuroImage 17, 825–841 (2002).

62. Smith, S. M. Fast robust automated brain extraction. Hum. Brain Mapp. 17, 143–155 (2002).

63. Avants, B. B., Tustison, N. & Song, G. Advanced normalization tools (ANTS). Insight J. 2, 1–35 (2009).

64. Fonov, V., Evans, A., McKinstry, R., Almli, C. & Collins, D. Unbiased nonlinear average age-appropriate brain templates from birth to adulthood. NeuroImage 47, S102 (2009).

65. Fonov, V. et al. Unbiased average age-appropriate atlases for pediatric studies. NeuroImage 54, 313–327 (2011).

66. O’Doherty, J. P., Buchanan, T. W., Seymour, B. & Dolan, R. J. Predictive Neural Coding of Reward Preference Involves Dissociable Responses in Human Ventral Midbrain and Ventral Striatum. Neuron 49, 157–166 (2006).

67. Sitek, K. R. et al. Mapping the human subcortical auditory system using histology, postmortem MRI and in vivo MRI at 7T. eLife 8, 1–36 (2019).

